# Characterization of m^6^A modifiers and RNA modifications in uterine fibroids

**DOI:** 10.1101/2023.08.07.552278

**Authors:** Jitu W. George, Rosa A. Cancino, Jennifer L. Griffin Miller, Fang Qiu, Qishan Lin, M Jordan Rowley, Varghese M. Chennathukuzhi, John S. Davis

## Abstract

Uterine leiomyoma or fibroids are the most common prevalent noncancerous tumors of the uterine muscle layer. Common symptoms associated with fibroids include pelvic pain, heavy menstrual bleeding, anemia, and pelvic pressure. These tumors are a leading cause of gynecological care but lack long-term therapy as the origin and development of fibroids are not well understood. Several next-generation sequencing technologies have been performed to identify the underlying genetic and epigenetic basis of fibroids. However, there remains a systemic gap in our understanding of molecular and biological process that define uterine fibroids. Recent epitranscriptomics studies have unraveled RNA modifications that are associated with all forms of RNA and are thought to influence both normal physiological functions and the progression of diseases. We quantified RNA expression profiles by analyzing publicly available RNA-seq data for 15 known epigenetic mediators to identify their expression profile in uterine fibroids compared to myometrium. To validate our findings, we performed RT-qPCR on a separate cohort of uterine fibroids targeting these modifiers confirming our RNA-seq data. We then examined protein profiles of key m^6^A modifiers in fibroids and their matched myometrium. In concordance with our RNA expression profiles, no significant differences were observed in these proteins in uterine fibroids compared to myometrium. To determine abundance of RNA modifications, mRNA and small RNA from fibroids and matched myometrium were analyzed by UHPLC MS/MS. In addition to the prevalent N6-methyladenosine (m^6^A), we identified 11 other known modifiers but did not identify any aberrant expression in fibroids. We then mined a previously published dataset and identified differential expression of m^6^A modifiers that were specific to fibroid genetic sub-type. Our analysis also identified m^6^A consensus motifs on genes previously identified to be dysregulated in uterine fibroids. Overall, using state-of-the-art mass spectrometry, RNA expression and protein profiles, we characterized and identified differentially expressed m^6^A modifiers in relation to driver mutations. Despite the use of several different approaches, we identified limited differential expression of RNA modifiers and associated modifications in uterine fibroids. However, considering the highly heterogenous genomic and cellular nature of fibroids, and the possible contribution of single molecule m^6^A modifications to fibroid pathology, there is a need for greater in-depth characterization of m^6^A marks and modifiers in a larger and varied patient cohort.

## Introduction

Uterine fibroids are steroid hormone-responsive, benign neoplasms of the uterus composed of smooth muscles, fibroblasts, and an abundance of extracellular matrix (1, 2). These benign tumors are estimated to occur in around 70% of women and clinically manifest in 30% of women by the age of 50 (3). Common clinical symptoms associated with fibroids are heavy bleeding, pain, infertility, and recurrent pregnancy loss (4, 5). Nonsteroidal anti-inflammatory drugs (NSAIDs), gonadotropin-releasing hormone agonists, elective estrogen receptor modulators, aromatase inhibitors, anti-progestins, and progesterone receptor modulators provide some relief but are only recommended for short-term use due to variable side effects and disease recurrence. (2). Accordingly, the lack of long-term therapeutic options and clinical morbidity associated with fibroids, hysterectomy remains the only option for many women (6). The cost of fibroid treatment and related health complications cost to the U.S. health care system is annually estimated to cost $5.9 to $34.4 billion (4, 7). Fibroids remain a significant burden on both health care costs and quality of life.

Several factors, including age, parity, ethnicity, and family history, are thought to act as drivers, but direct evidence identifying etiology of the disease has been difficult (4). Genome wide exome sequencing identified that 30-90% of fibroids, depending on patient ethnicity and fibroid number, contained mutations in the second exon of the mediator complex subunit 12 (*MED12*) gene (8). Additionally, chromosomal rearrangement at 12q15 and 6q21, leading to overexpression of the High Mobility Group A1/2 (*HMGA1/2*) gene, has been observed in 8-35% of fibroids (9, 10). Biallelic inactivation of *FH* (Fumarate Hydratase), deletion of the collagen genes *COL4A5* and *COL4A6*, and mutations of the SNF2-Related CBP Activator Protein (SRCAP) complex subunits are among the rarer subtypes that have been reported in uterine fibroids (11, 12). These chromosomal events trigger sub-type specific gene expression patterns that are either shared or unique to the genetic event (12, 13).

Since its discovery in 1974, methylation of adenosine on RNA (m^6^A or N^6^-methyladenosine) has emerged a major post-transcriptional RNA modification (14). The past decade has seen a resurgence in examining the role of m^6^A RNA modifications in regulating RNA processing, splicing, export, stability, and translation (15). Transcriptome-wide profiling of m^6^A modification identified the modification as wide-spread, highly selective, and dynamic in nature, with levels varying in development and cellular stress (16, 17). Addition of m^6^A sites is catalyzed by the “writer” proteins, specifically by the catalytic activity of Methyltransferase-like protein 3, METTL3, and target RNA binding activity of methyltransferase-like protein 14 (METTL14) (18). In addition to METTL3, other regulatory proteins involved in the process include methyltransferase-like protein 16 (METTL16), Wilms tumor 1-associated protein (WTAP), RNA-binding motif 15 (RBM15), Cbl proto-oncogene-like protein 1 (CBLL1), zinc-finger CCCH-type-containing 13 (ZC3H13), and Vir-like m^6^A methyltransferase-associated (VIRMA) (or also known as KIAA1429) (18, 19). Following addition of m^6^A modification, biological consequence is regulated by specific RNA-binding proteins, or “readers”. These readers recognize and bind to the DRACH (D=A, G or U, R= G or A, and H= A, C, or U) consensus sequence of modified RNA transcripts leading to regulation of gene expression and modulation of diverse processes including splicing, mRNA stabilization, and translation efficiency. Reader proteins include YTHD domain protein family (YTHDC1, YTHDC2, YTHDF1, YTHD2, and YTHDF3) (20). In addition, heterogenous nuclear ribonucleoprotein (HNRNP) protein, HNRNPA2B1, has been shown to regulate m^6^A-modified transcript including a subset of primary miRNA (21). Two proteins, fat mass and obesity-associated protein (FTO) and alkB homologue 5 (ALKBH5) have been identified as m^6^A demethylases or “erasers” due to their ability to remove m^6^A marks (20). Altered expression of these eraser proteins can contribute to atypical cellular functions and physiological activity thereby promoting tumorigenesis (18, 20).

To date multiple studies carried out by us, and others have identified differential DNA methylation patterns, histone modifications, altered miRNA and long noncoding RNA expression as they relate to uterine fibroidogenesis (11–13, 22–24). However, an in-depth characterization of m^6^A modifiers and RNA modifications in uterine fibroids is lacking. The goal of this study was to investigate expression patterns of the vast array of m^6^A modifier proteins and RNA modifications in both mRNA and small RNA as they relate to uterine fibroids compared to myometrium.

## Material and methods

### Human Tissue collection and Sample Preparation

Matched samples of human myometrium and fibroid samples were collected from pre-menopausal women undergoing hysterectomy for symptomatic uterine fibroids. Use of human tissue was approved by the University of Nebraska Medical Center (IRB# 112-21-EP) and University of Kansas Medical Center (IRB#: 5929), and all patients signed a written informed consent form to donate tissue for this study. Human samples were processed as previously described (13). Upon arrival, samples were minced and sub-divided for a) RNA extraction and b) protein isolation for western blots and then immediately flash frozen and stored at −140 °C.

### RNA extraction

Total RNA was extracted from freshly frozen samples as previously described (13). Following total RNA extraction by Trizol, mRNA from fibroids and matched myometrium (n=6) was isolated by two rounds of purification using olido-dT Dynabeads mRNA DIRECT Micro kit (Dynabeads) according to manufacturer’s protocol. Depending on patient sample, 8-50 µg total RNA was used per purification column. Integrity and purity of isolated mRNA was evaluated using Fragment Analyzer Automated CE System (Advanced Analytical Technologies, Inc). RNA Integrity Numbers (RINs) were used to evaluate integrity and samples with RIN >7.0 were considered intact and used for further downstream LC-MS/MS analysis.

Small RNA (<200 bp) was isolated from fibroids and matched myometrium (n= 5) using mirVana miRNA isolation kit (Thermo Fisher Scientific) according to manufacturer’s protocol. Following isolation, small RNA purity was confirmed using Fragment Analyzer Automated CE System (Advanced Analytical Technologies, Inc).

### Quantitative real-time PCR (RT-qPCR)

cDNA was synthesized from 1µg total RNA using qScript cDNA synthesis kit (Quantbio, Beverly, MA). Quantitative Real rime PCR (RT-qPCR) analysis was performed on genes of interest (**Supplementary Table 1**) using SYBRGreen (BioRad). Sso Fast EvaGreen Supermix was performed to analyze gene expression on a BioRad CFX96 Real-Time System (BioRad, Hercules, CA). Relative quantification of gene of interest was established using RPL17 as reference and calculated using the comparative Ct method.

### LC-MS/MS Analysis of RNA Chemical Modifications

Measurement of the levels of RNA chemical modifications was performed using ultra-performance liquid chromatography coupled with tandem mass spectrometry (UHPLC-MS/MS) by a method similar as described (25–27). Briefly, Total amount of 100 ng of small RNA or mRNA was digested with a Nucleoside Digestion Mix (New England BioLabs) according to the manufacturer’s instruction. The digested samples were then lyophilized and reconstituted in 100 µl of RNAse-free water, 0.01% formic acid prior to UHPLC-MS/MS analysis. The UHPLC-MS/MS analysis was accomplished on a Waters XEVO TQ-S^TM^ (Waters Corporation, USA) triple quadruple tandem mass spectrometer equipped with an electrospray source (ESI) source maintained at 150 °C and a capillary voltage of 1 kV. Nitrogen was used as the nebulizer gas, which was maintained at 7 bars pressure, flow rate of 1000 l/h and at temperature of 500°C. UHPLC-MS/MS analysis was performed in ESI positive-ion mode using multiple-reaction monitoring (MRM) from ion transitions previously individually determined for these RNA chemical modifications (28). A Waters ACQUITY UPLC^TM^ HSS T3 guard column, 2.1x 5 mm, 1.8 µm, attached to a HSS T3 column, 2.1 x50 mm, 1.7 µm was used for the separation. Mobile phases included RNAse-free water (18 MΩcm^-1^) containing 0.01% formic acid (Buffer A) and 50% acetonitrile (v/v) in Buffer A (Buffer B). The digested nucleotides were eluted at a flow rate of 0.2 ml/min with a gradient as follows: 0-1 min, 0 %B; ramp to 0.2% B in 1.4 min; then to 0.8% in 1.4 min, 3.8-5.2 min, 0.8-1.8% B; 5.2-6.6 min, 1.8-3.2%B; 6.6-10 min, 3.2-5.0% B;10-13.5 min, 5-8%B; 13.5-18 min, 8-30%B; in 0.5 min to 100% B and kept for 1.5 min. The total run time was 25 min. The column oven temperature was kept at 25 °C and the sample injection volume was 10 µl. Three injections were performed for each sample. Data acquisition and analysis were performed with MassLynx V4.1 and TargetLynx. Calibration curves were plotted using linear regression with a weight factor of 1/x.

### Western Blot

Protein was isolated from fibroids and matched myometrium by homogenization in Radioimmunoprecipitation assay (RIPA) buffer supplemented with protease and phosphatase inhibitors. Following isolation, lysate protein concentration was quantified using the Pierce BCA Protein assay kit and 10 µg of protein were separated on a 10% SDS-PAGE and transferred onto nitrocellulose membranes (Amersham). Membranes were blocked in 5% BSA in Tris-buffered saline with 0.1% Tween-20 (TBST) at room temperature for 1 h and probed with METTL3 (1:1000; 15073-1-AP; Proteintech), METTL14 (1:1000; 26158-1-AP; Proteintech), RBM15 (1:1000; VIRMA (1:1000; 25712-1-AP; Proteintech), WTAP (1:1000; 10200-1-AP; Proteintech), CBLL1 (1:1000; 21179-1-AP; Proteintech), FTO (1:1000; 27226-1-AP; Proteintech), ALKBH5 (11:1000; 6837-1-AP; Proteintech), and β-actin (1:5000; A5441; Sigma) antibodies at 4 °C overnight. Following washes in TBST, membranes were blocked in secondary HRP-conjugated antibodies (1:10,000) in 5% BSA in TBST for 1 h, washed, and imaged using iBright system (ThermoFisher). Densitometry analysis was performed using Image J. All protein levels were normalized to respective ACTB which served as loading control.

### Statistics

RNA-seq data (13) were filtered, normalized, and converted to log2-counts per million (CPM) value per sample. The study included two factors: within-subject factor tissue (Fibroid or Normal) and the between-subject factor race (B or W). For each gene, linear mixed models were used to assess the race effect and tissue effect on the mean normalized gene expression levels (log_2_CPM), accounting for correlations between observations from the same patient. We are interested in the following hypothesis testing: whether the race effect on the normalized gene expression level was significant in fibroids and normal tissues respectively, whether the tissue effect on the normalized gene expression level is significant in black and white patients respectively, and whether the tissue effect alters in black patients compared to white patients per gene. The associated p-values for each comparison on each gene were adjusted by the false discovery rate (FDR) method of Benjamini and Hochberg method (1) due to multiple hypotheses testing. Differentially expressed genes were identified as those having an FDR below 0.05.

Statistical analyses were performed using GraphPad Prism 9.0. The Student’s t-test was used to compare fibroid samples to myometrium and significance level was set at P< 0.05. Sample numbers are indicated in all figure legends. Data presented in graphs are expressed as mean ± SEM.

## Results

### Transcriptomic expression of m^6^A regulators in uterine fibroids

To assess a possible role for m^6^A modifications in uterine fibroids, we first tested whether m^6^A regulators are differentially expressed in fibroids versus normal myometrium. Paired analysis of published RNA-seq (13) data revealed little to no difference in the transcriptomic expression levels of writers (*METTL3, METTL14, METTL4, CBLL1, VIRMA, WTAP, RBM15, ZC3H13*), readers (*HNRNPA2, YTHDF1, YTHDF2, YTHDC1, YTHDC2*), and erasers (*FTO* and *ALKBH5*) in normal myometrium and fibroids [(13)] (**Figure 1a**). Analysis of a separate published microarray dataset (29) confirmed the overall lack of difference (**Figure 1b**). While most genes tested displayed little or no overall differences (**Suppl. Figure 1**), we did note one candidate (*RBM15*) that displayed a slight, but statistically significant difference when comparing the transcript levels across the samples (**Figure 1c**). To confirm our finding, we performed RT-qPCR on a separate set of fibroids and matched myometrium patient samples, however, for most m^6^A modifiers (*METTL3*, *YTHDC1*, *FTO*) we did not see correlation with RNA-seq data probably due to the extreme modest changes observed between fibroids and myometrium (**Suppl. Figure 2**). RT-qPCR confirmed *RBM15* expression was slightly, but significantly different when comparing fibroids with matched myometrium (**Figure 1d**).

**Figure 1:**
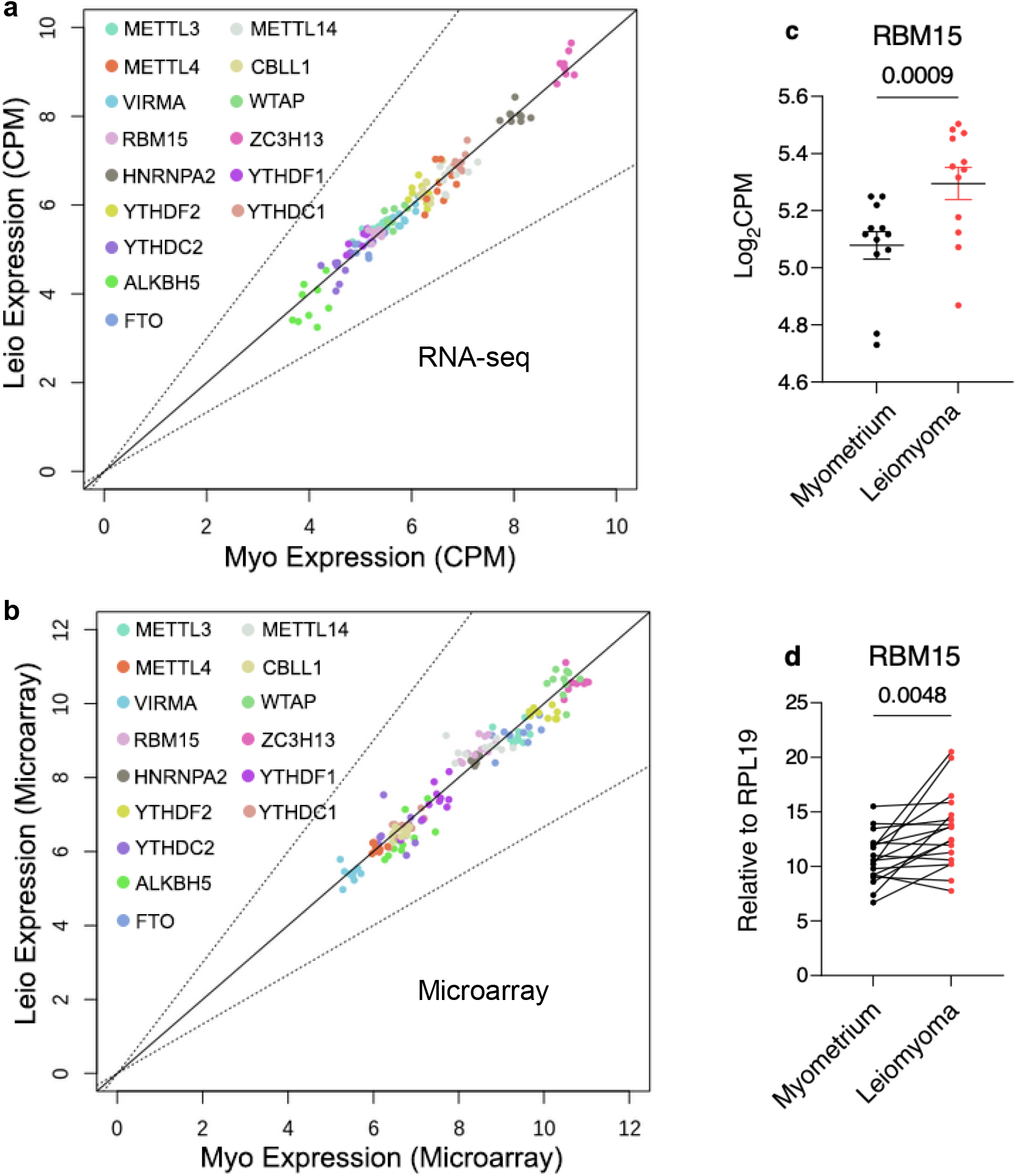
Transcriptomic analysis of m^6^A modifiers in uterine fibroids. **a-b.** RNA-seq (a) and microarray (b) analysis of leiomyoma and matched myometrium. Transcript abundance quantified as counts per million (CPM) from myometrium (Myo, x-axis) and leiomyomas (Leio, y-axis). Diagonal represents no differences, while the dashed lines represent 1.5 fold changes. **c**. The log2 counts per million (log_2_ CPM) from myometrium (n=9) and fibroids (n=12) of *RBM15*. Data are represented as means ± SEM. Statistically significant differences between groups were calculated with paired student’s t-test. *P*-values for each comparison is reported. **d**. Relative expression of *RBM15* in myometrium and matched fibroid samples measured by RT-qPCR (n=18). Results are presented relative to *RPL17*. Statistically significant differences between groups were calculated with paired student’s t-test. *P*-values for each comparison is reported.

Previous studies identified differential expression of mRNA and miRNA between fibroids isolated from Black and White women indicating these candidate factors could drive racial disparity of the disease (30, 31). We therefore examined published RNA-seq data for racial differences in the expression of m^6^A modifiers (13, 32, 33). Overall, we detected higher variation among datapoints (**Suppl. Figure 3**) and did not detect race specific molecular disparity within normal myometrium or fibroids, within Black and White women (**Suppl. Figure 4**). However, while *RBM15* was statistically significantly upregulated in fibroids compared to normal myometria in White women (**Figure 2a**, p=0.035), it showed similar trends but was not statistically significant in Black women (**Figure 2b**, p=0.06).

**Figure 2:**
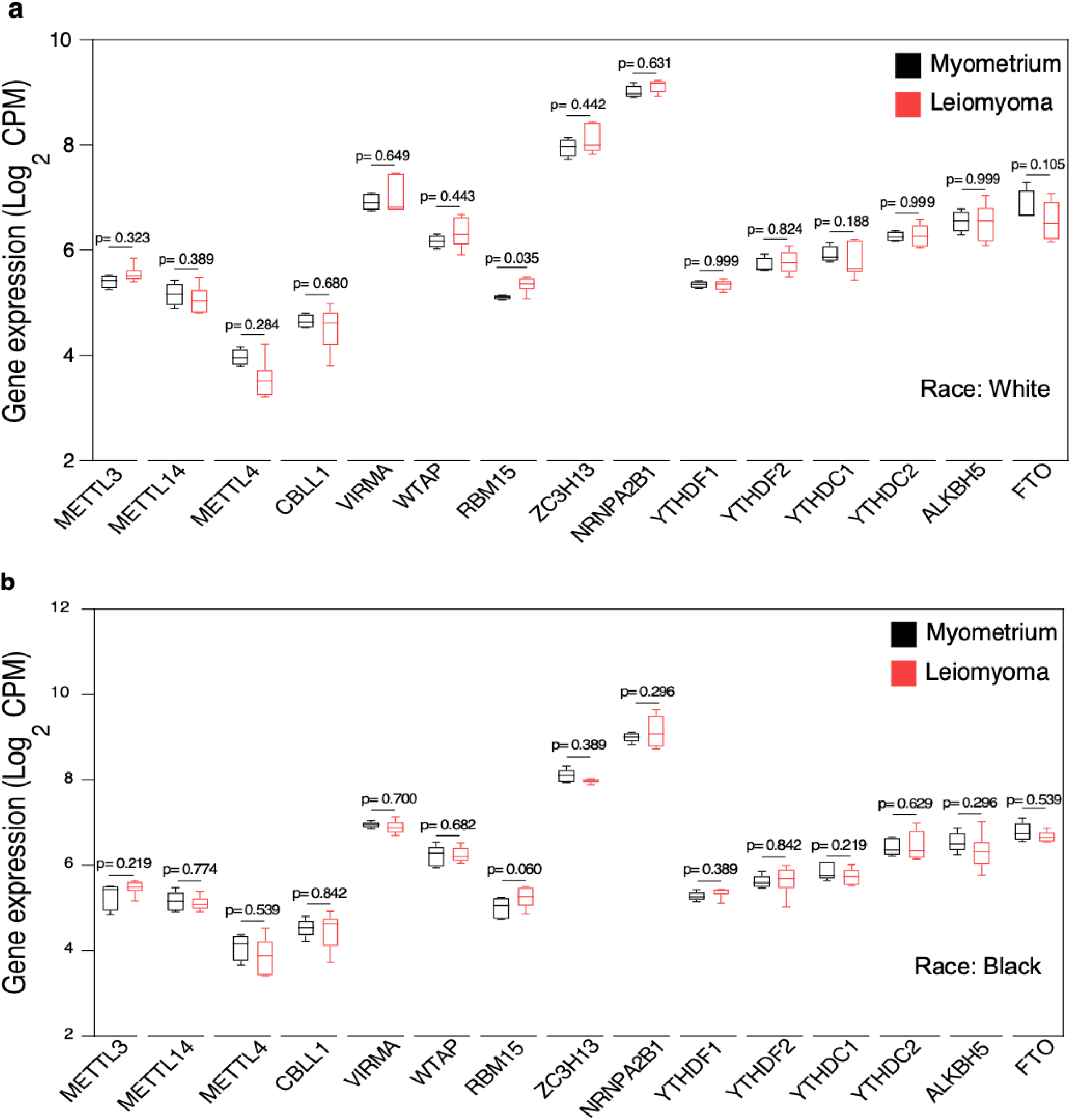
Transcriptomic analysis of m^6^A modifiers in leiomyoma and matched myometrium in White and Black women. RNA-seq analysis (SRP166862) of normalized expression of leiomyoma and matched myometrium (GSE120854). The y-axis is signal abundance quantified as log2 counts per million (log2 CPM). **a.** mRNA expression in normal myometrium (n=4) and leiomyoma (n=6) from white women. **b.** mRNA expression in normal myometrium (n=5) and leiomyoma (n=6) from black women. Data are represented as means ± SEM. *FDR* for each comparison is reported.

### Protein expression of m^6^A modifiers in uterine fibroids

To identify differences in protein levels, we performed western blot analysis on key m^6^A modifier proteins (**Figure 3**). We found that, while some individuals displayed differences (**Figure 3a**), the overall levels of these m^6^A modifiers were not significantly different between normal myometrium (n= 19) and fibroids (n= 26) (**Figure 3b**). Indeed, even though our expression analysis revealed subtle differences in RBM15 transcript levels (**Figure 1c-d**), there was no significant difference in RBM15 protein expression within fibroids (n=23) and matched myometrium (n=19) (**Figure 3a-b)**. Altogether, these data indicate an overall lack of protein expression differences of m^6^A modifiers between fibroids and myometrium.

**Figure 3:**
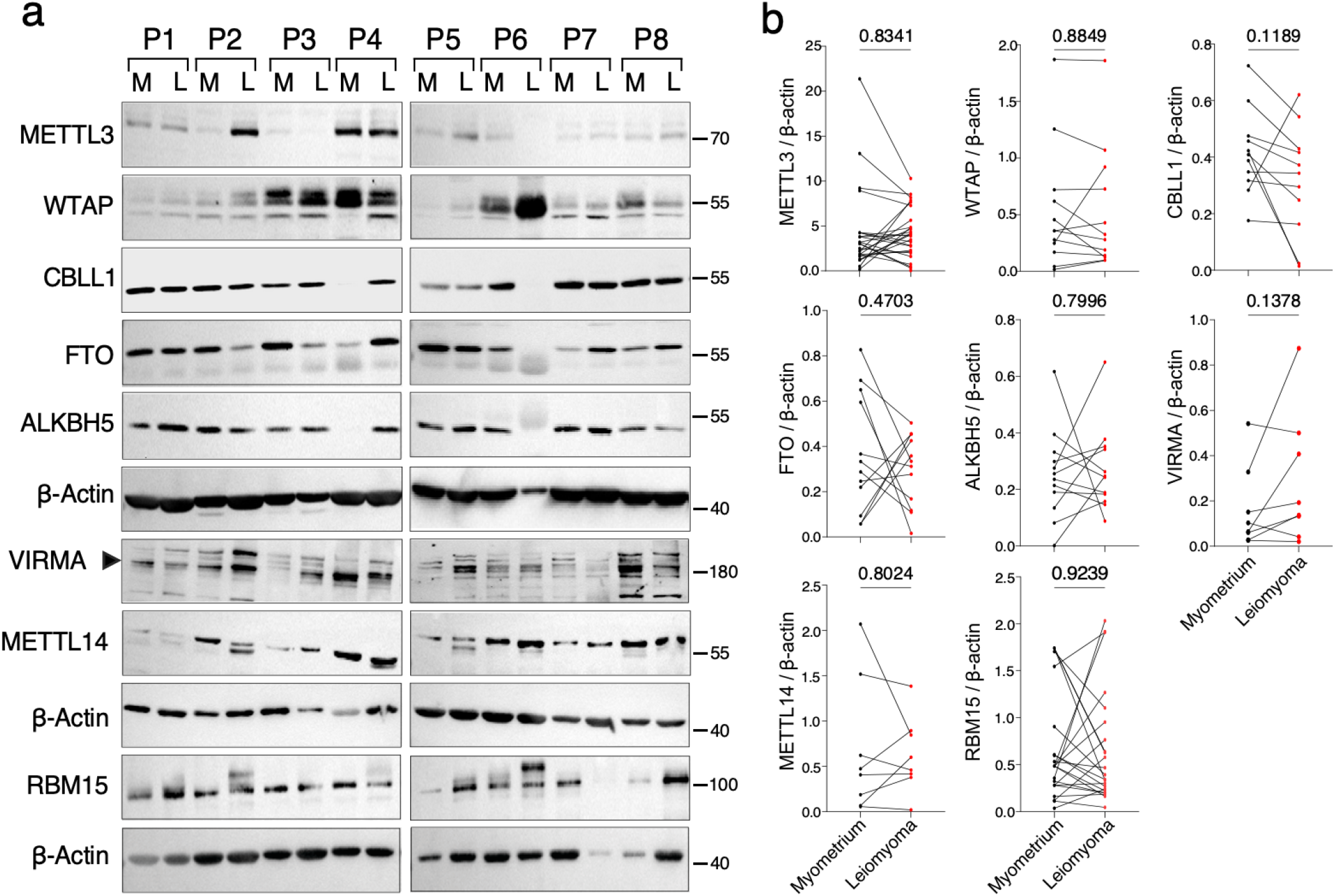
Western blot analysis of m^6^A modifiers in leiomyoma. Proteins were isolated from uterine leiomyoma and matched myometrium and probed for m^6^A modifier proteins. (**A**) Representative western blots of METTL3, WTAP, CBLL1, VIRMA, METTL14, RBM15, FTO, and ALKBH5. ACTB was used as loading control. M=Myometrium; L=Leiomyoma. P denotes individual patient samples. (**B**) Data was quantified for METTL3 (myometrium (n=19) and leiomyoma (n=26), RBM15 ( myometrium (n=15) and leiomyoma (n=23), WTAP, CBLL1, VIRMA, METTL14, FTO, and ALKBH5, were quantified from 12 paired fibroids and matched myometrium patient samples. Statistically significant differences between groups were calculated with paired student’s t-test. *P*-values for each comparison is reported.

### Abundance of mRNA and small RNA modifications in uterine fibroids

In light of the overall lack of changes in m^6^A modifier expression, we next considered the possibility of differential methylase and demethylase activity between fibroids and myometrium. To measure m^6^A, mRNA was isolated, and concentration of modified RNA nucleosides was measured by UHPLC-MS/MS. No differential signal abundance of m^6^A was observed between fibroids and matched myometrium (**Figure 4a**). We extended our analysis to other well characterized mRNA modifications that have been identified in eukaryotes and known to impact various cellular functions (34) to determine if these were dysregulated in uterine fibroids. We did not identify any additional modifications (m^5^C, m^7^G, ac^4^c, m^1^A, f^5^C, dA, ho^5^u) to have altered abundance in our fibroid samples versus myometrium samples (**Figure 4b-h**).

**Figure 4:**
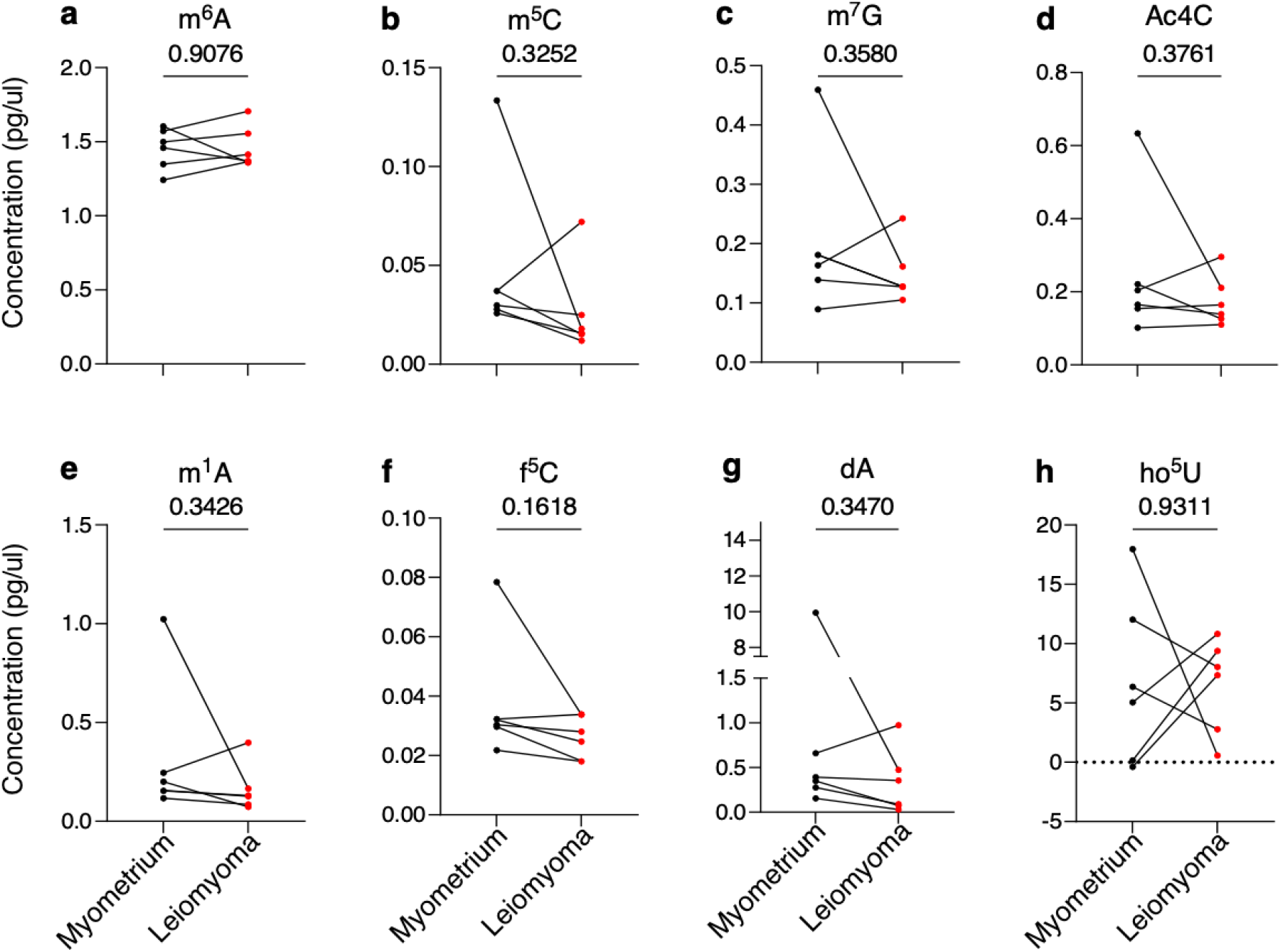
mRNA modifications from normal myometrium and matched fibroids. The y-axis is signal abundance quantified as pg/µl myometrium (n=6) and matched fibroids (n=6). Changes in (**a**) N^6^-methyladenosine (m^6^A), (**b**) 5-methylycytosine (m^5^C), (**c**) N^7^-methylguanosine (m^7^G), (**d**) N^4^-acetylcytidine (ac^4^C), (**e**) N^1^-methyladenosine (m^1^A), (**f**) 5-Formylcytidine (f^5^C), (**g**) 2’-deoxyadenosine (dA), (**h**) 5-hydroxyuridine (ho^5^U) were measured. Statistically significant differences between groups were calculated with paired student’s t-test. *P*-values for each comparison is reported.

In addition to mRNA, small RNAs (<200 bps) are known to harbor diverse RNA modifications that can modulate complex biological processes (35). Small RNAs have been identified to be differentially expressed in fibroids and thought to regulate multiple processes that influence uterine fibroid development and progression (31). Following isolation of small RNA from normal myometrium and matched fibroids, we measured post translational RNA modifications. We were unable to detect differential expression of m^6^A levels in our fibroid samples and myometrium samples (**Suppl. Fig 5a**). We simultaneously measured other modified nucleoside abundance including, m^1^A, i^6^A, ac^4^c, m^5^C, m^3^C, f^5^C, m^1^G, m^7^G, mo^5^U, ho^5^U, m^5^U that have been implicated to regulate translational machinery and influence physiological processes (35). Our analysis did not identify these modifications to be differentially expressed (**Suppl. Fig 5b-l**), indicating an absence of preferential small RNA modifications in fibroids.

### Characterization of m6A modifiers with respect to genetic sub-type

Uterine fibroids are driven by multiple driver mutations, including *MED12*, *HMGA1*, *HMGA2*, *FH* and the more recently characterized mutation in the SRCAP complex subunits (11–13). Multiple studies both from our lab and others have shown that normal myometrium and these driver mutations form separate transcription clusters indicating altered pathways are activated in these genetic subtypes (11–13). To define a broader clinical perspective, we mined gene expression profiles of 162 normal myometrium and 190 fibroid samples that were recently published by Berta et al (11). These fibroid samples were divided into *MED12* (n=38), *HMGA2* (n=44), *HMGA1* (n=62), *FH* (n=15), *YEATS* (n=16) and *OM* (n=15) allowing capture of majority of fibroid subtypes as described (11). We mapped each genetic sub-type against fold change to determine if these modifiers were preferentially expressed (**Figure 5)**. We identified statistical, though modest changes in majority of epigenetic regulators based on mutation status. Among the readers, *METTL3* was found to be significantly upregulated in *HMGA2* (log2-fold 0.11) and *YEATS* (log 2-fold 0.15) sub-type fibroids (**Figure 5a**). *METTL14* on the other hand was significantly downregulated in *MED12* (log 2-fold - 0.16) and *HMGA2* (log 2-fold −0.08) fibroids, while upregulated in *YEATS* (log 2-fold 0.12) fibroids but was not statistically significant (**Figure 5b**). In congruence with our RNA-seq analysis (**Figure 1, Suppl. Figure 1**), *RBM15* was upregulated in almost all fibroid sub-types (**Figure 5g**). Among writer proteins, *YTHDF1* was upregulated in majority of fibroid sub-types (**Figure 5k**), while *YTHDC2* was significantly downregulated in *HMGA1* (log 2-fold −0.09), *FH* (log 2-fold −0.16), *YEATS* (log 2-fold - 0.13) fibroid subtype (**Figure 5n**). There was elevated expression of m^6^A demethylase *ALKBH5* in majority of fibroid sub-types (**Figure 5o**), while mRNA expression of *FTO* was significantly decreased in all sub-types (**Figure 5p**).

**Figure 5:**
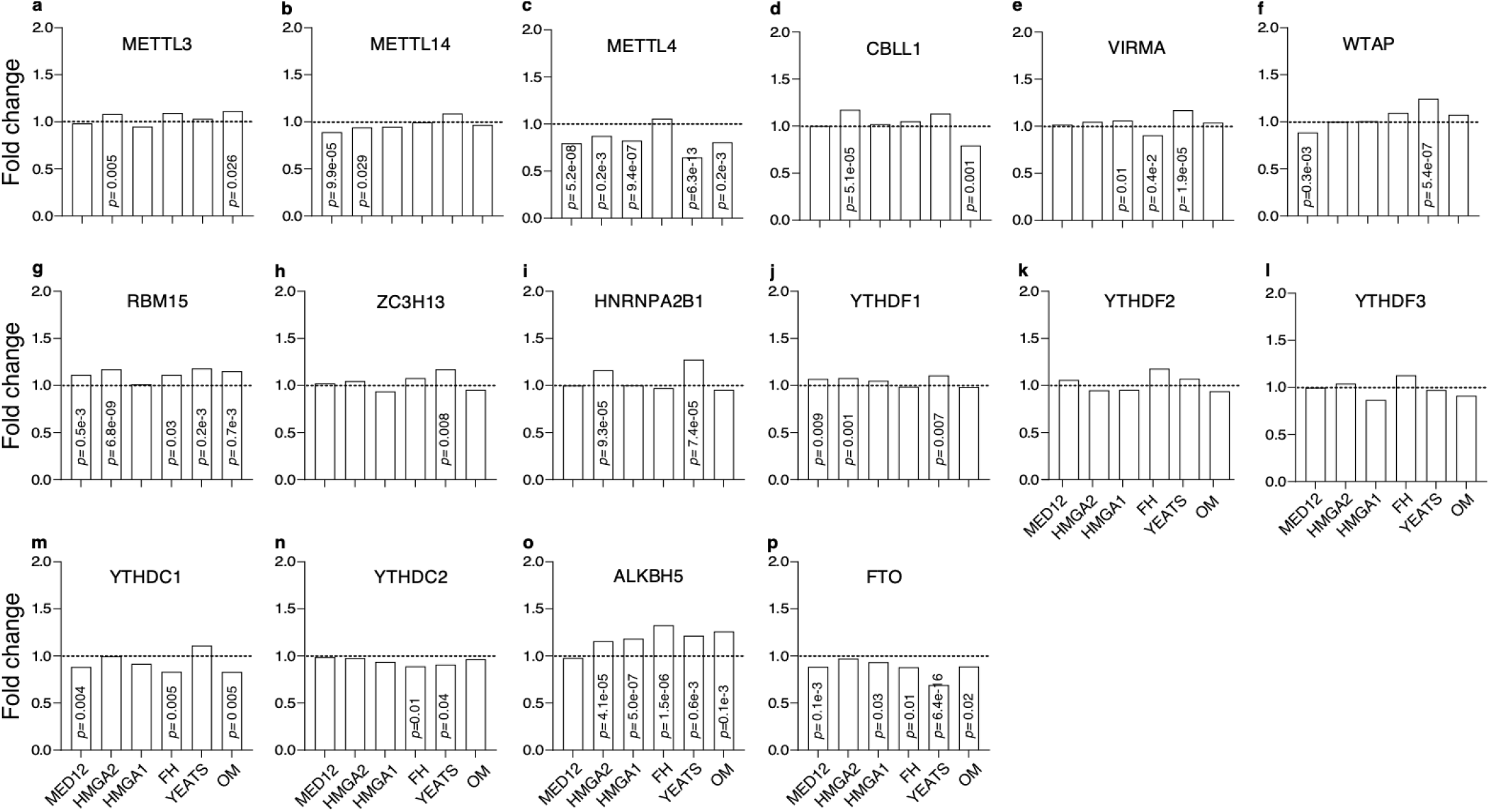
Expression profile of m^6^A modifiers in relation to fibroid genetic sub-types. Gene expression of m^6^A modifiers mapped in relation to fibroid genetic sub-types as defined by Berta et al (11). Y-axis denotes fold-change, and each genetic subtype is denoted in the X-axis. *MED12* (Mediator complex subunit 12), *HMGA1/2* (High Mobility group A1/2), *FH* (Fumarate Hydratase), *YEATS* (YEATS domain containing 4), *OM* (Other Mutations as defined by alteration of other members of the SRCAP complex subunits). *MED12* (n=38), *HMGA2* (n=44), *HMGA1* (n=62), *FH* (n=15), *YEATS* (n=16) and *OM* (n=15). Reader proteins (**a-h**), Writers (**i-n**), and Erasers (**o, p**). Statistical significance was set at FDR <0.05 and value for each comparison is reported.

We next harnessed mammalian m^6^A predictor, SRAMP (<u>s</u>equence-based <u>R</u>NA <u>a</u>denosine <u>m</u>ethylation site <u>p</u>redictor)(36) to predict if key fibroid genes identified by us and others (11–13), had m^6^A sites in their mRNA (**Supplementary Table 2**). We identified a collection of putative m^6^A sites with varying levels of confidence in steroid hormone receptor genes, (Progesterone Receptor (*PGR)*, Estrogen Receptor (*ESR1*)), transcription factors (Pleomorphic adenoma gene 1 (*PLAG1),* Pappalysin 2, (*PAPPA2)*, Chromobox 2 (*CBX2)*, Chromobox 4 (*CBX4)*, Chromobox 8 (*CBX8)*, SATB homeobox 2 *(SATB2)*), DNA repair protein (RAD51 Paralog B (*RAD51B)*), steroidogenic genes (Hydroxysteroid 17-beta dehydrogenase 6 (*HSD17B6)*, steroid 5 alpha-reductase 2 (*SRD5A2)*, tryptophan 2,3-dioxygenase (*TDO2)*), collagen associated genes (ADAM metallopeptidase domain 12 (*ADAM12)*, Collagen type I alpha 1 chain (*COL1A1)*, collagen type 3 alpha 1 chain (*COL3A1)*, Periostin (*POSTN)*) and growth factors Cyclin D1 (*CCND1),* vascular endothelial growth factor A *(VEGFA*)). These in silico analysis suggests that key fibroid genes are possibly susceptible to RNA modification and subsequent transcriptional regulation. However, in-depth analysis will be required to define transcriptome-wide m^6^A location and efficacy of these marks in fibroid etiology.

## Discussion

Over 170 RNA modifications have been identified of which m^6^A accounts for the most abundant and widespread mRNA internal modification (20, 37). The diverse distribution patterns, its dynamic nature, and ability to regulate multiple physiological processes has added another layer to post-transcriptional regulation. Multiple studies have now identified that regulation of m^6^A is driven by methyltransferases, demethylases, or reader proteins and dysregulation of which is closely associated with human cancers (38). In reproductive cancers, m^6^A modifiers were identified to regulate ovarian, endometrial, and cervical cancer (39, 40). However, characterization of m^6^A modification proteins have not been defined in uterine fibroids. Here, for the first time, we provide an in-depth characterization of major modifiers of m^6^A modification as it relates to uterine fibroids.

The m^6^A methyltransferase complex is comprised of a METTL3/METTL14 heterodimer core that adds m^6^A in a highly specific manner (41). Due to the enzymatic ability of METTL3 which allows addition of m^6^A to nuclear RNA, we paid special attention to both its transcriptomic and protein abundance in fibroids. Our analysis did not identify differential expression in either RNA or protein of METTL3 in uterine fibroids in absence of mutation status. Lack of differential m^6^A RNA modification was also observed in our LC-MS/MS data from both purified fractions of mRNA and small RNA (**Figure 3A**; Supplementary Figure 3A). However, increased expression of METTL3 has been reported and could be attributed to patient and fibroid heterogeneity (42, 43). When broken down by fibroid sub-type, we see mild increased expression of METTL3 mRNA in *HMGA2* and *YEATS* fibroid (**Figure 4A**). METTL3 and METTL14 form a 1:1 heterodimer and recognize the DRACH motif leading to induction of m^6^A modification on mRNA (44, 45). Inactivation or deletion of METTL14 results in depletion of m^6^A in mRNA, identifying it as a core regulator of m^6^A addition (46). While no significant changes were observed in transcriptomic and protein levels of METTL14 in global fibroid samples, we identified changes when pared down by fibroid genetic sub-types. In addition to METTL3 and METTL14, other core components are known to mediate m^6^A addition. Among these include WTAP which interacts and anchors METTL3 and METTL14 to nuclear speckles regulating gene expression and alternative splicing (34). WTAP interacts with another well-known m^6^A mediator, VIRMA (46, 47). Apart from VIRMA, ZC3H13 is another WTAP interactor and shown to be required for nuclear localization of the writer complex (20, 37). CBLL1 has also been shown to couple with WTAP (20, 37) and loss of CBLL1 led to reduction of global m^6^A levels, identifying it as another writer protein. WTAP expression was found to be increased in *YEATS* fibroid and decreased in *MED12* mutants. While VIRMA and ZC3H13 were increased in *YEATS* fibroid and CBLL1 was increased only in *HMGA2* fibroids. Among writer proteins, we saw increased transcriptomic expression of RBM15 in both global (**Figure 1G**), by race (**Suppl Figure 2 a**) in multiple fibroid genetic sub-type (**Figure 4G**). However, protein levels were not significantly changed in fibroids (**Figure 2a, 2b**). RBM15 has been identified as part of m^6^A writer complex and shown to bind the long non-coding RNA, *XIST*. Knocking down RBM15 decreased m^6^A methylation on *XIST* RNA, leading to reduced *XIST* mediated gene silencing (48). *XIST* has been identified to regulate fibroid pathology by sponging miR-29c and miR-200c leading to increased expression of COL1A1, COL3A1, and FN1, key regulators of extracellular accumulation (49). As an RNA binding protein, in addition to its role as a m^6^A modulator, RBM15 regulates splicing of key differentiation genes involved in hematopoietic stem cells quiescence (50). A hypothesis put forward to define uterine fibroid etiology is reprogramming of myometrial stem cells leading to fibroid development (51). Whether RBM15 plays a role in regulating cell fate decision of myometrial cells and if so, does it mediate its action through m^6^A or alternative splicing, and its contribution to uterine fibroid pathogenesis remains to be explored.

There are five YTH domain-containing proteins (YTHDC1-2 and YTHDF1-3) that have been structurally identified to recognize m^6^A through a conserved aromatic cage. YTHDF 1-3 is cytoplasmic, YTHDC1 is predominantly nuclear, while YTHDC2 can be both nuclear and cytoplasmic. YTHDF1 has been linked to enhanced translation of m^6^A mRNA, while YTHDF2 binding leads to RNA degradation bought about by recruitment of CCR4-NOT deadenylation complex. Finally, YTHDF3 cooperatively binds to YTHDF1 and YTHDF2 regulating translation and degradation thereby impacting gene expression profile of m^6^A-containing mRNA (20, 52). Among YTHDF1-3, statistical significance was identified only in YTHDF1 (**Figure 4K-O**). In addition to regulating m^6^A mRNA, YTHDC1 also appears to mediate function of long noncoding RNA, in particularly *XIST* and regulating transcriptional silencing of the X-chromosome (48, 53). Transcript levels of both YTHDC1 and YTHDC2 were found to decreased in some variation in all fibroid genetic subtypes.

Two m^6^A demethylases, FTO and ALKBH5, have been identified that are able to convert m^6^A to A and regulate global m^6^A levels. FTO and ALKBH5 have been reported to be dysregulated in diverse diseases leading to m^6^A demethylation, modulation of gene expression downstream and influencing biological consequence (54, 55). With relation to fibroid genetic subtype, we saw an inverse correlation with regards to transcript profiles between ALKBH5 and FTO, indicating that these demethylases are nonredundant and exhibit distinct epigenetic regulation.

There is a strong racial disparity in the disease, with Black women presenting with an earlier onset of the disease and greater severity (56). We postulated that fibroids from Black women could exhibit differential expression of m^6^A modifiers when compared to White women. Transcriptomic analysis of m^6^A modifiers identified increased transcript expression of RBM15 in fibroids from White women (**Suppl Figure 2**), but not Black (**Suppl Figure 2b**) indicating that there might be differential m^6^A levels between fibroids obtained from White and Black women but will need confirmation in a much larger cohort of patient samples.

Altered levels of modifiers based on genetic sub-type, led us to explore if genes known to be associated with fibroids had specific m^6^A methylation patterns. As uterine fibroids rely on estrogen and progesterone to grow, increased levels of ESR1 and PGR may affect underlying molecular pathways driving fibroid growth and progression. Traditionally anti-progestins are prescribed to reduce uterine bleeding and decrease fibroid volume (57–62). In silico analysis identified multiples sites on *ESR1* and *PGR* that could harbor m^6^A modification with moderate and high confidence, indicating that transcript abundance of these key steroidogenic receptor molecules may be regulated post-transcriptionally. SRAMP analysis also predicted m^6^A modification sites on several transcription factors that were implicated to regulate uterine fibroids, namely, *PLAG1*, *PAPPA2*, *CBX2*, *CBX4*, *CBX8*, *SATB2*. Transcript levels of *PLAG1* and *PAPPA2* were identified to have elevated expression in HMGA1/2 uterine fibroids (13). While *CBX2*, *CBX4*, *CBX8*, *SATB2*, have been previously implicated in uterine fibroidogenesis(11, 13, 63). DNA repair protein, RAD51B is upregulated in *MED12*, *HMGA1*, and *HMGA2* fibroids suggesting a cell response to genomic instability and a possible “second-hit” pushing normal myometrial cells to tumorigeneses (64). In addition, in silico analysis identified m^6^A marks on TDO2 mRNA, a key enzyme catalyzing the conversion of tryptophan to kynurenine which was recently identified to be upregulated in *MED12* mutant fibroids and dependent on race (65–67). TDO2 inhibitor, 680C91, reduced expression of *COL1A1* and *COL3A1*, genes involved in collagen production and extracellular matrix (ECM) accumulation, in primary uterine fibroid culture (65). More recently, primary myometrial cells treated with mono(2-ethyl-5-hydroxyhexyl) phthalate (MEHHP), increased expression of *TDO2*, promoting tryptophan metabolism. Depletion of TDO2 reduced proliferative action of MEHHP on primary fibroid cells identifying it as pro-survival factor. Our in-silico analysis also identified m^6^A marks on *COL1A1*, *COL3A1*, and *POSTN*, known structural constituents of ECM organization. Notably, in triple negative breast cancer cells, increased expression of METTL3 was negatively correlated with COL3A1 expression (68). We and others identified increased expression of POSTN and characterized its role as potential regulator of fibroidogenesis (69, 70). POSTN has also been identified to be regulated by through m^6^A modification during cardiac remodeling (71). *CCND1* and *VEGFA*, other known regulators of uterine fibroids (13, 72, 73) were similarly ordained with m^6^A modifications in their RNA. CCND1 has been identified to be regulated through its m^6^A modification and influence hematopoietic stem/progenitor cells differentiation (74).

In summary, while global transcriptomics and protein levels were unchanged, we identified modest genetic sub-type expression of m^6^A modifiers. Our analysis did identify transcript levels of *RBM15* to be consistently increased but differential expression in protein levels were not detected. While modest (1-1.4-fold), we identified statistically significant differential expression, indicating that driver mutations could regulate m^6^A deposition in a fibroid specific subtype manner. Characterization of key fibroid genes identified multiple m^6^A marks indicating the possibility of interplay between methylation and mRNA expression and downstream deregulation of biological processes. However, in-depth sequencing and characterization of m^6^A sites will be needed to be performed to further define the possibility of m^6^A in fibroid pathology. Since protein abundance can be post-transcriptionally regulated and protein levels are not always correlated with mRNA levels (75), the discovery of m^6^A marks and modifiers in uterine fibroids opens the field to additional effector molecules that were previously unappreciated in tumor formation. While our studies did not identify difference in protein expression or RNA modifications in uterine fibroids, an in-depth approach with larger patient cohorts with regards to genetic sub-types and race will be needed to define expression profiles and validate possible influence of RNA modifications on fibroid pathogenesis and progression.

## Acknowledgments

The authors have no conflict to declare.

---

**Supplementary Figure 1:**
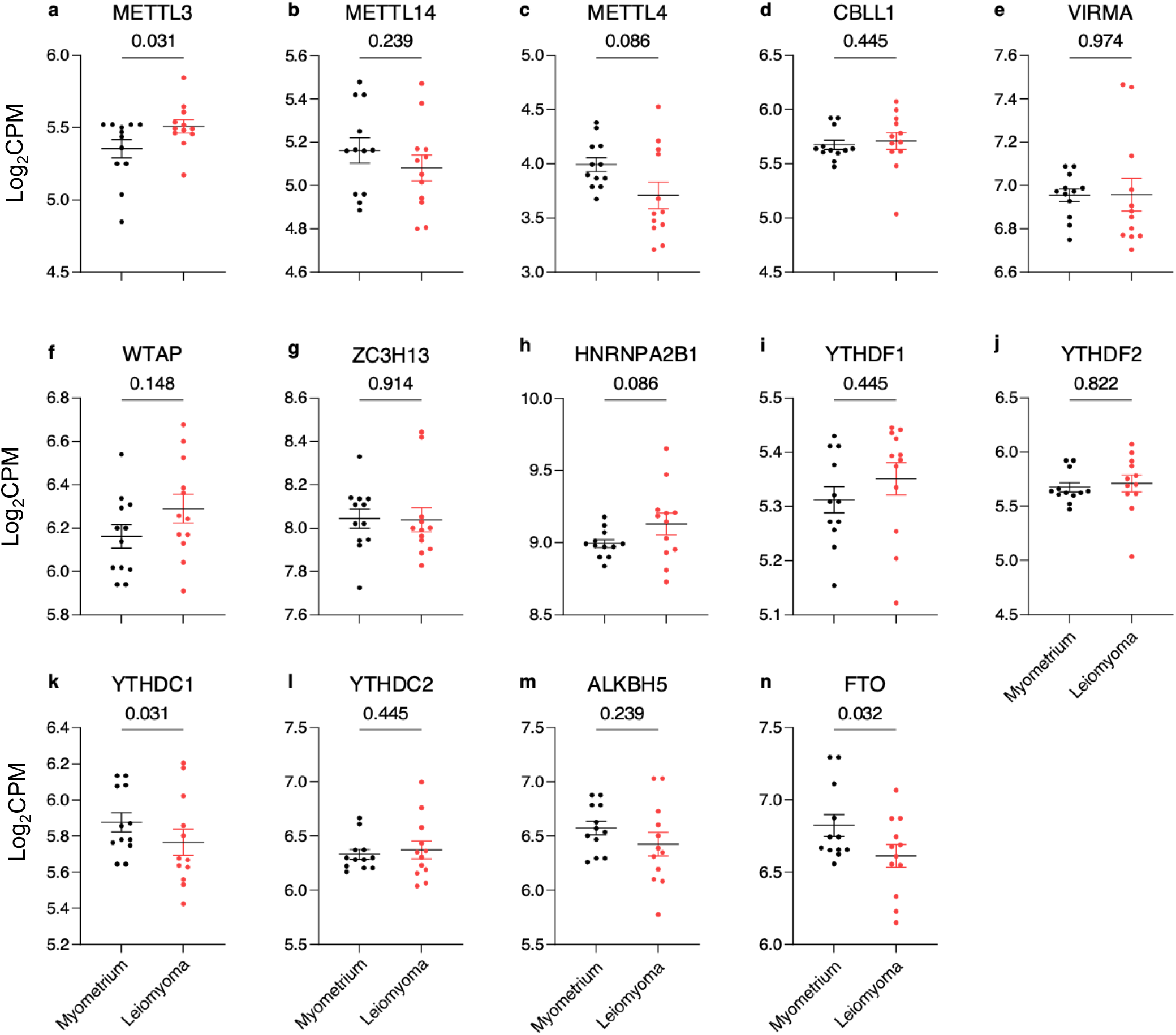
Transcriptomic analysis of m^6^A modifiers in uterine fibroids. RNA-seq analysis of normalized expression of leiomyoma and matched myometrium (SRP166862). The y-axis is signal abundance quantified as log2 counts per million (log_2_CPM) from myometrium (n=9) and fibroids (n=12). **a-f**. mRNA expression of m6A writers (*METTL3*, *METTL14*, *METTL4*, *CBLL1*, *VIRMA*, *WTAP*). **g-l**. Readers (*ZC3H13*, *HNRNPA2*, *YTHDF1*, *YTHDF2*, *YTHDC1*, *YTHDC2*), and erasers (**m, n**), (*ALKBH5*, *FTO*). Data are represented as means ± SEM. Statistically significant differences between groups were calculated with paired student’s t-test. *FDR* values for each comparison is reported.

**Supplementary Figure 2:**
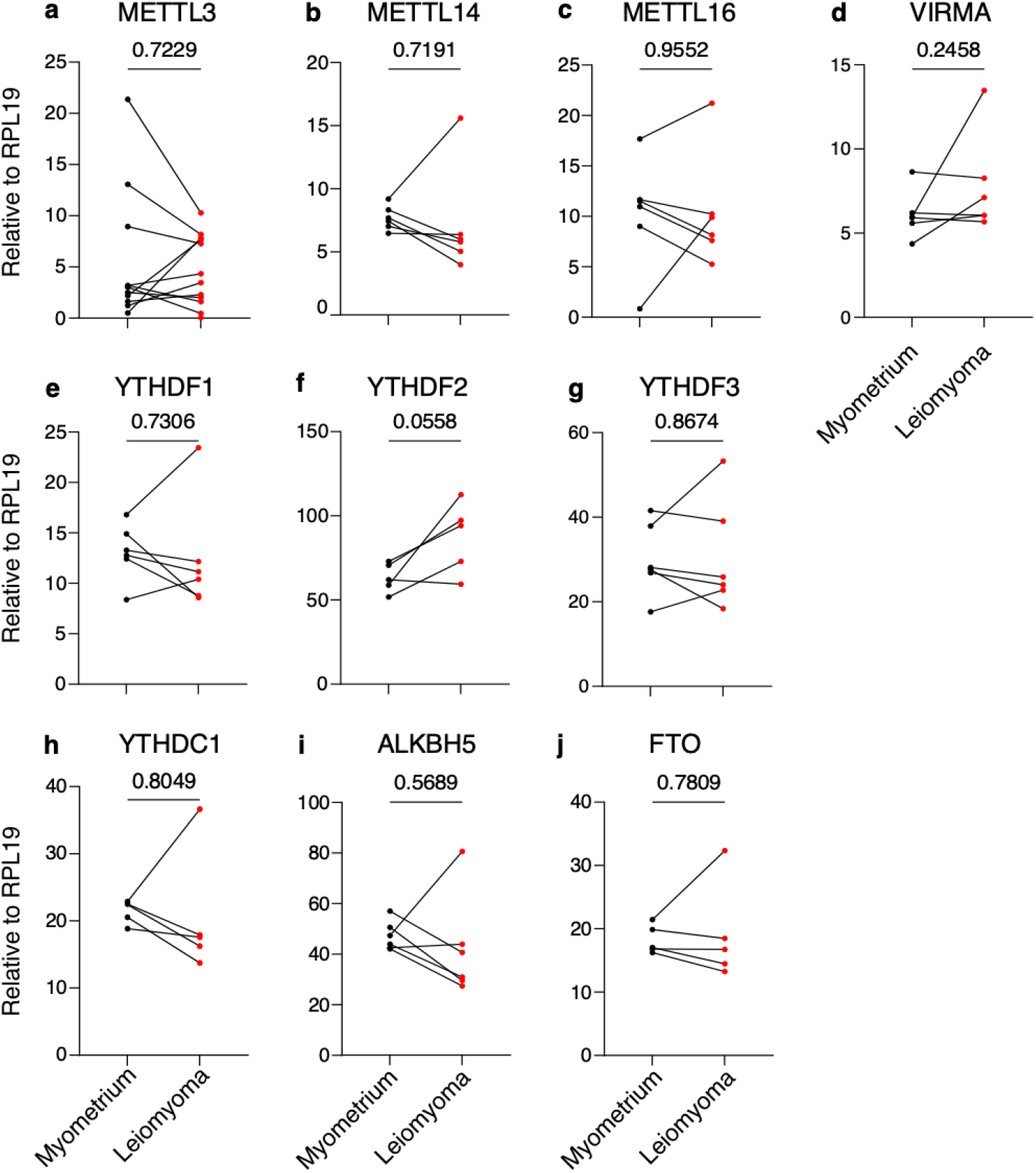
RT-qPCR analysis of m^6^A modifiers in uterine fibroids and matched myometrium. RNA levels of (**a**) *METTL3* (n=12), (**b**) *METTL14* (n=6), (**c**) *METTL16* (n=6), (**d**) *VIRMA* (n=6), (**e**) *YTHDF1* (n=6), (**f**) *YTHDF2* (n=6), (**g**) *YTHDF3* (n=6), (**h**) *YTHDC1* (n=6), (**i**) *ALKBH5* (n=6), (**j**) *FTO* (n=6). Results are presented relative to *RPL17*. Statistically significant differences between groups were calculated with paired student’s t-test. *P*-values for each comparison is reported.

**Supplementary Figure 3:**
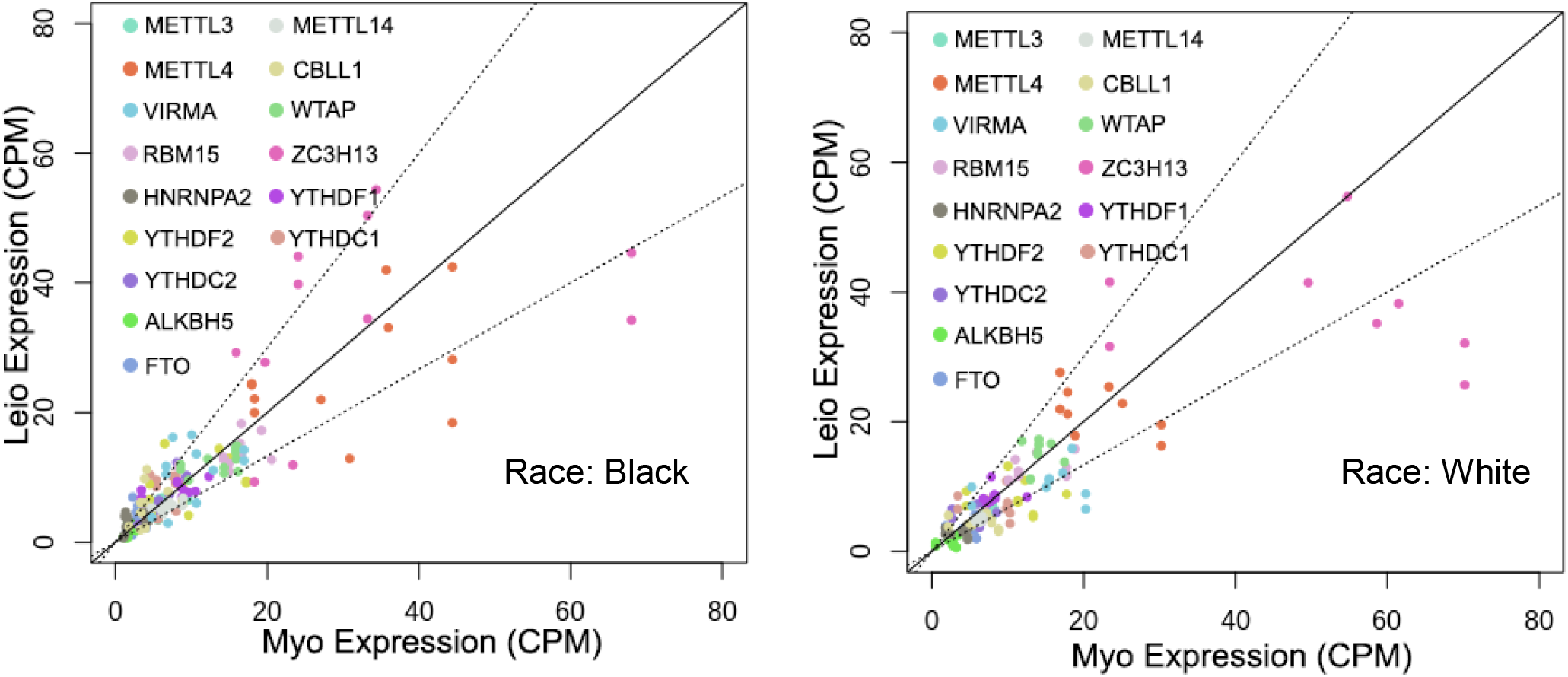
Analysis of leiomyoma and matched myometrium (PRJNA859428, GSE207209. Transcript abundance quantified as counts per million (CPM) from myometrium (Myo, x-axis) and leiomyomas (Leio, y-axis). Diagonal represents no differences, while the dashed lines represent 1.5 fold changes.

**Supplementary Figure 4:**
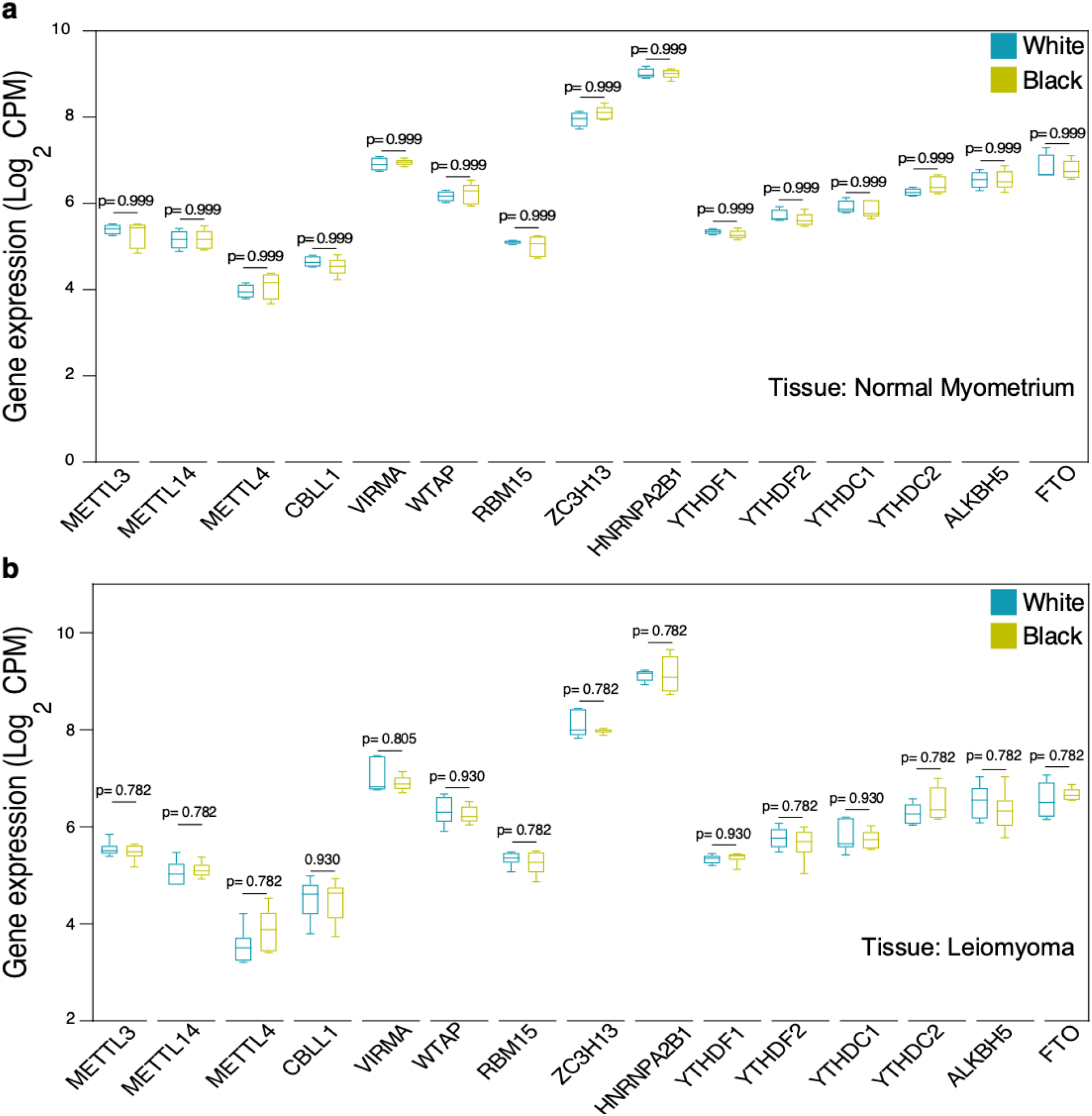
Transcriptomic analysis of m^6^A modifiers in normal myometrium and leiomyoma in White and Black women. RNA-seq analysis (SRP166862) of normalized expression of leiomyoma and matched myometrium (GSE120854). The y-axis is signal abundance quantified as log2 counts per million (log2 CPM). **a**. mRNA expression in normal myometrium from White (n=4) and Black (n=5) women. **b**. mRNA expression in leiomyoma from White (n=6) and Black (n=6). Data are represented as means ± SEM. *FDR* for each comparison is reported.

**Supplementary Figure 5:**
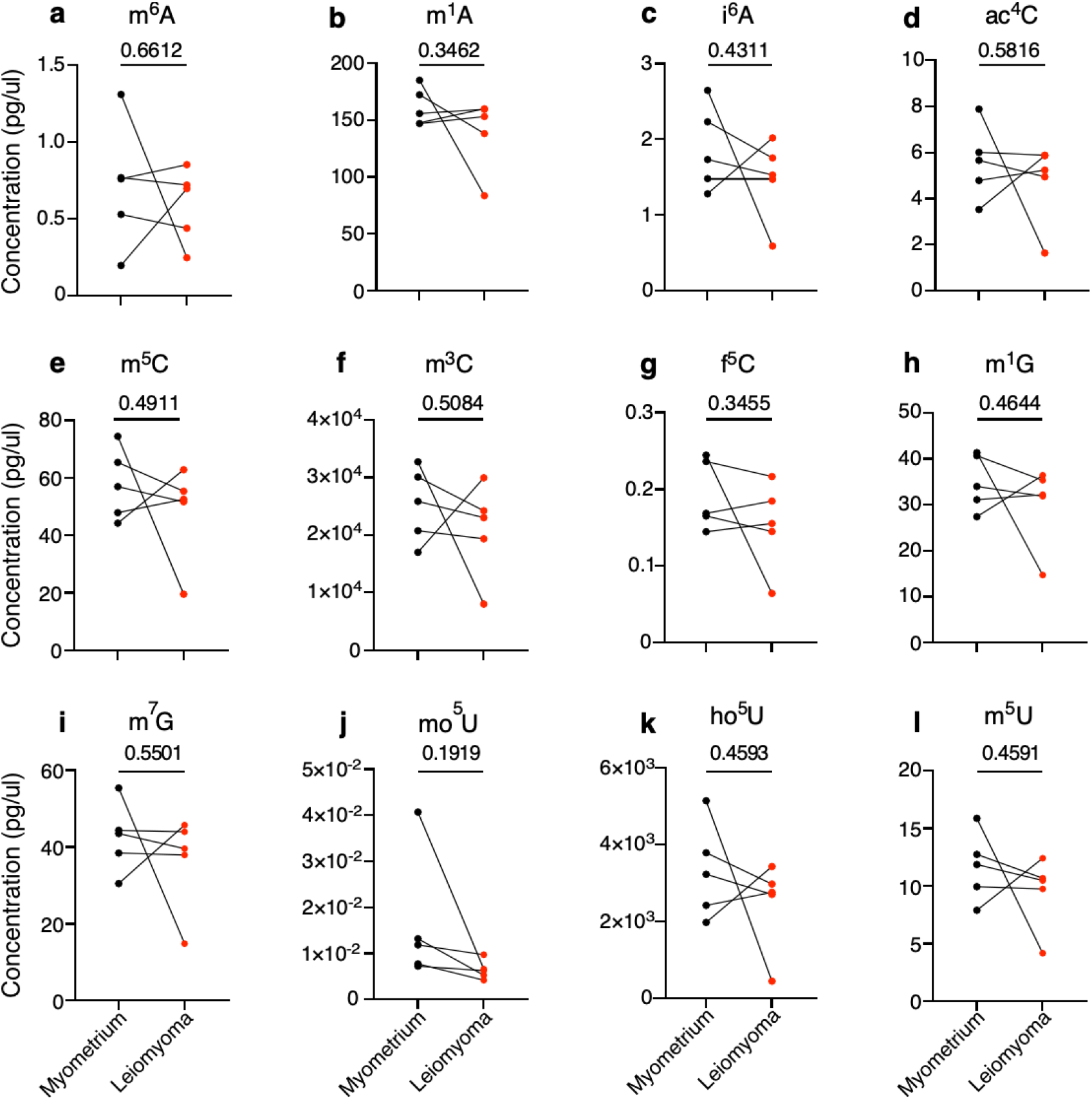
small RNA modifications from normal myometrium and matched fibroids. The y-axis is signal abundance quantified as pg/µl myometrium (n=5) and matched fibroids (n=5). Changes in (**a**) N^6^-methyladenosine (m^6^A), (**b**) N^1^-methyladenosine (m^1^A), (**c**) N^6^-isopentenyladenosine (i6A), (**d**) N^4^-acetylcytidine (ac^4^C), (**e**) 5-methylycytosine (m^5^C), (**f**) N^3^-methylcytidine (m^3^C), (**g**) 5-Formylcytidine (f^5^C), (**h**) N^1^ methylguanosine (m^1^G), (**i**) N^7^-methylguanosine (m^7^G), (**j**) 5-methoxyuridine (mo^5^U) (**k**) 5-hydroxyuridine (ho^5^U), (**l**) 5-methyluridine (m^5^U) were measured. Statistically significant differences between groups were calculated with paired student’s t-test. *P*-values for each comparison is reported.

